# Testing the reliability of AI-generated protein structures

**DOI:** 10.64898/2026.06.11.731682

**Authors:** Amanda Xu, Steven L. Salzberg

## Abstract

Although AlphaFold2 and its competitors have demonstrated remarkable abilities to predict protein structure, more work is needed to explore the limitations of these methods. Here we investigated the reliability of AlphaFold2 and ColabFold by creating a set of realistic but false protein sequences, using ColabFold to predict their structure, and then asking how often the program produces a high-scoring structure for a sequence that does not represent a protein. We determined that AlphaFold2 has a very small but non-zero false positive rate, estimated here at approximately 1 in 435 if one uses a threshold pLDDT score of 70 to define positive predictions. We also discovered, serendipitously, that some high-scoring sequences in the human genome were not false positives, but instead were previously unknown and un-annotated pseudogenes. These latter findings indicate that some well-established human annotations of protein-coding genes may have incorrectly extended the 5’ untranslated regions too far. They also suggest that AlphaFold2’s false positive rate is low enough that almost any high-scoring structure, even in a noncoding region, is worthy of further investigation.

## Introduction

AlphaFold2 [1] was a major step forward in the prediction of protein structures from sequence alone. It was the first computational method that could reliably predict structures whose accuracy rivaled that of experimental methods, and as a result it was hailed as the Breakthrough of the Year in 2021 [2], and in 2024 the senior authors of the AlphaFold2 paper shared the Nobel Prize in chemistry.

AlphaFold2 (AF2) and its free online version ColabFold have been used to fold millions of proteins in the years since their first release, including every protein in the human genome [3]. The newer AlphaFold3 (AF3) system can predict the structure of interacting proteins, albeit with somewhat less accuracy than when predicting isolated structures [4]. Work from our lab has shown that when considering a set of isoforms (or splice variants) of the same protein, AF2’s scores can be used as accurate indicators of which protein isoforms are likely to be functional [5].

And yet despite the proven track record of AF2 at predicting structures, few efforts have focused on measuring its false positive rate. Or to put this question differently, how often will AF2 predict a “good”protein structure for a sequence that does not actually fold into that structure? The extreme complexity of the deep learning model embedded within the program makes it impossible to answer this question analytically. In addition, given the difficulty of conventional laboratory methods, it is simply not feasible to synthesize predicted protein sequences and try to solve their structures experimentally. Thus there is no easy way to check that a high-scoring structure produced by AF2 or AF3 is correct.

Both AF2 and ColabFold generate per-residue confidence scores for each structure they produce, which are known as pLDDT (predicted Local Distance Difference Test) scores and which range from 0 to 100. The average pLDDT across an entire structure is generally used to estimate the structure’s accuracy, with numbers above 70 considered to be generally correct (or high-confidence), and numbers above 90 considered to represent highly accurate and stable structures [1].

Two previous studies did consider variations on the question of AF2’s false positive rate. The first study [6] looked at AntiFam, a database of hidden Markov models (HMMs) representing families of proteins that were annotated incorrectly; i.e., they are translations of open reading frames (ORFs) found in actual DNA sequences that do *not* represent functional proteins [7]. In the Monzon *et al*. study [6], the authors used AF2 to fold all 250 models in AntiFam, and found that a large majority of AntiFam proteins scored poorly, with only one entry longer than 100 amino acids receiving a pLDDT score above 70. The one exception, which received a score >90, turned out to be a genuinely functional protein, which they subsequently removed from AntiFam, and which now is represented in GenBank (accession ANF00096.1) as a bona fide protein, with assigned function as a subunit of cytochrome oxidase. Thus in this prior study, AlphaFold2 assigned pLDDT scores below 70 to all “false” proteins translated from ORFs in actual genomic DNA. In that same study, the investigators used AF2 to score randomly-generated amino acid sequences of lengths varying from 10 to 200aa, and found that 100% of random sequences >80aa had pLDDT scores below 50, with scores growing smaller as lengths increased.

A second recent study looked at the propensity of AF2 to assign high scores to artificially-generated proteins that comprised a series of 10 perfect repeats, where the repeat unit ranged from 5 to 30 amino acids in length [8]. That study found that both AF2 and AF3 tended to create β-solenoid structures from these artificial repeat-containing proteins, but that the structures were often biophysically implausible. AF2 gave high pLDDT scores (>=70) to a majority of the structures for these artificial proteins, while AF3 tended to assign much lower scores, although it produced similar β-solenoid structures.

Thus one might conclude from the AntiFam study that the false positive rate for sequences with pLDDT > 70 is very low, perhaps close to zero, although the Pratt *et al*. study [8] showed that high scores might be assigned to artificially constructed sequences such as proteins comprised entirely of short repeating elements. In the experiments described below, we attempted to estimate AF2’s false positive rate empirically using a much larger number of examples taken from real genomes, but using protein sequences that, *a priori*, were extremely unlikely to form stable structures. Our goal here was to explore the use of AF2 as a tool for predicting the structures of amino acid sequences translated from ORFs that appear in real genomes but that are not currently annotated as protein-coding genes, and to ask whether a high pLDDT score might be a strong indicator that the sequence represented a gene.

## Results

We collected all of the annotated protein-coding genes from the human genome and from four bacterial genomes: *Bacillus anthracis, Helicobacter pylori, Salmonella enterica*, and *Staphylococcus aureus*. For human, we used the MANE annotation database [9], a curated set of very high-quality annotations that contains only one splice variant (or isoform) for each protein-coding gene. For each gene in each species, we identified the longest ORF on the opposite strand of the annotated gene, under the assumption that DNA virtually never encodes a protein in both directions. We then selected those opposite-strand ORFs (osORFs) that were at least 300 base pairs in length, in order to generate a set of “false” proteins of 100 amino acids or longer. This generated 1363 bacterial proteins and 7320 human proteins, all of which were then folded using ColabFold [10], a free version of AF2. We then extracted all predicted structures that had an average pLDDT score of at least 70 for further analysis. Note that none of the osORFs from *S. aureus* had scores >70, and therefore our discussion below describes translations from the other three bacteria.

Figure 1. shows a histogram of all scores for both bacterial and human false proteins translated from osORFs. The scores were distributed in an approximately bell-shaped curve with a peak around 40 for bacteria and 35 for human, as might be expected for non-functional amino acid sequences. However, on the upper end of the distributions, we identified a small number of proteins with scores ≥70 that were predicted to have stable structures.

**Figure 1.**
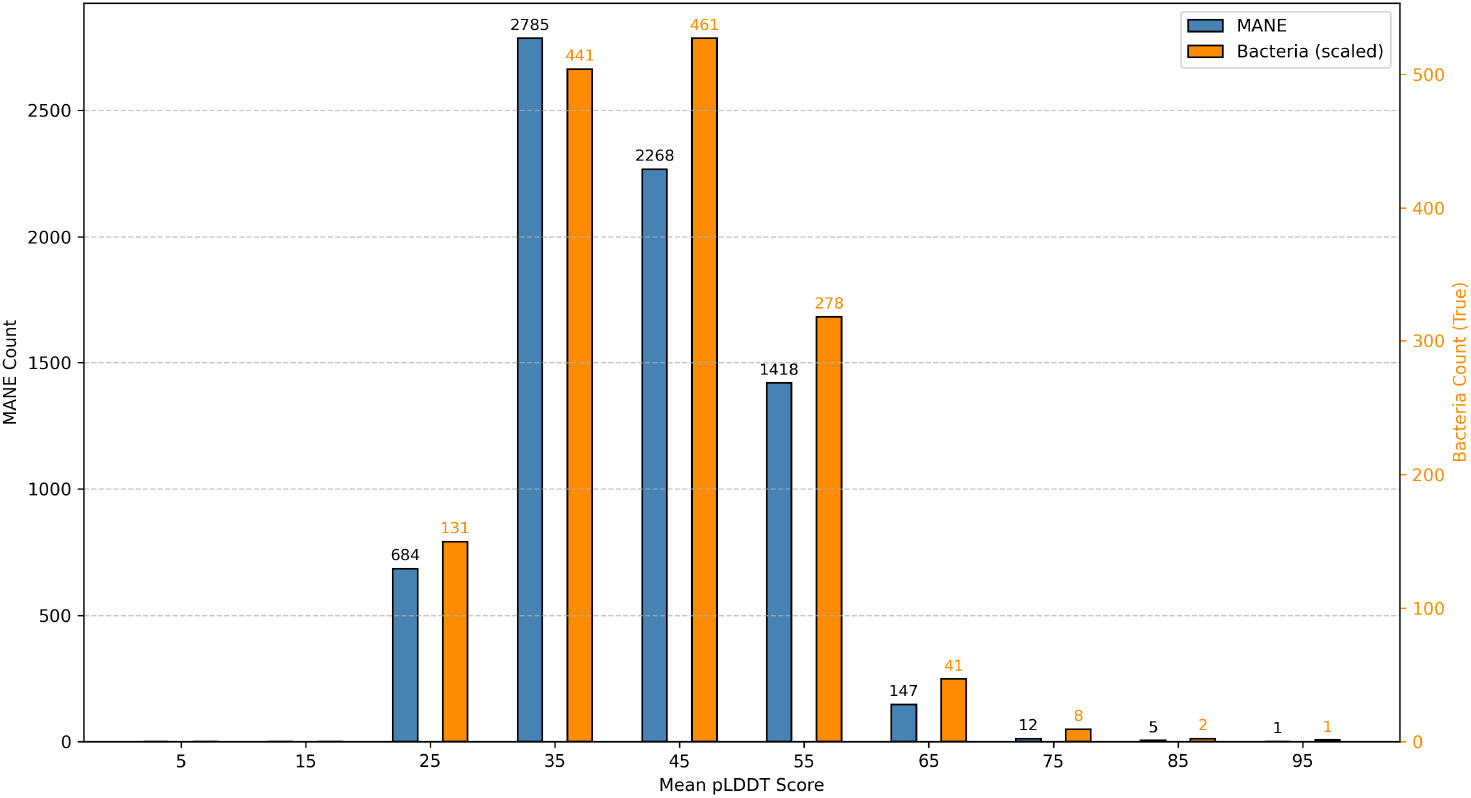
Mean pLDDT scores computed by ColabFold for protein sequences translated from osORFs in 7320 human protein-coding genes from the MANE annotation (blue, left axis) and in osORFs in 1363 proteins from three bacterial species (orange, right axis).

### High-scoring osORFs

Eighteen of the human osORF proteins and 11 of the bacterial osORF proteins received pLDDT scores above 70 (**Tables 1-3**). For all but one of the bacterial osORFs, the structures looked remarkably similar: they appeared to be simple alpha helices that spanned most of the sequence, often as two short helices separated by a kink in the structure. **Figure 2** illustrates four of these structures, found in the reverse strands of protein-coding genes in *B. anthracis* and *H. pylori*.

**Table 1.** Opposite-strand ORFs within human genes. This table lists opposite-strand open reading frames (osORFs) found within human protein-coding genes, where the osORF yielded an AlphaFold2 / ColabFold pLDDT score of 70 or greater. Shown are the primary human transcript ID (from RefSeq), the gene name, chromosome number, coordinates spanned by the transcript, the region where the osORF appears (CDS for protein-coding or 3UTR for the 3’ untranslated region), the coordinates of the osORF, the length in amino acids of the translated osORF, and its pLDDT score. Structures for these osORF proteins are shown in Suppl. Figure S1, labeled with the letters A-R shown in the first column.

| ID | Transcript ID | Gene | Chr | Transcript span | osORF region | osORF interval | Len (aa) | pLDDT |
| --- | --- | --- | --- | --- | --- | --- | --- | --- |
| A | NM_001304533.3 | NKAIN3 | chr8 | chr8:62248854–62984904 | 3UTR | chr8:62974140–62974595 | 151 | 90.47 |
| B | NM_002031.3 | FRK | chr6 | chr6:115931149–116060891 | 3UTR | chr6:115939708–115940115 | 135 | 87.05 |
| C | NM_030946.2 | OR14J1 | chr6 | chr6:29301701–29313017 | 3UTR | chr6:29310886–29311371 | 161 | 83.1 |
| D | NM_022117.4 | TSPYL2 | chrX | chrX:53082367–53088540 | CDS | chrX:53085713–53086048 | 111 | 81.9 |
| E | NM_021155.4 | CD209 | chr19 | chr19:7739993–7747534 | 3UTR | chr19:7740457–7740951 | 164 | 81.77 |
| F | NM_001013694.3 | SRRD | chr22 | chr22:26483877–26494658 | 3UTR | chr22:26492013–26492360 | 115 | 80.56 |
| G | NM_015299.3 | KHNYN | chr14 | chr14:24429954–24441843 | 3UTR | chr14:24440083–24440589 | 168 | 79.4 |
| H | NM_133462.4 | TTC14 | chr3 | chr3:180602163–180611130 | CDS | chr3:180607719–180607765;<br>180608701–180608810;<br>180609630–180609790 | 105 | 76.43 |
| I | NM_181741.4 | ORC4 | chr2 | chr2:147930396–148020737 | 3UTR | chr2:147931634–147932083 | 149 | 76.03 |
| J | NM_002275.4 | KRT15 | chr17 | chr17:41513745–41518890 | CDS | chr17:41518502–41518837 | 111 | 75.88 |
| K | NM_001396058.1 | OR2I1P | chr6 | chr6:29550407–29557773 | 3UTR | chr6:29555880–29556320 | 146 | 75.82 |
| L | NM_020347.4 | LZTFL1 | chr3 | chr3:45823316–45842119 | 3UTR | chr3:45824682–45825011 | 109 | 75.75 |
| M | NM_014594.3 | ZNF354C | chr5 | chr5:179060373–179083977 | 3UTR | chr5:179082673–179083083 | 136 | 74.66 |
| N | NM_080817.5 | GPR82 | chrX | chrX:41724181–41730130 | 3UTR | chrX:41729605–41729970 | 121 | 72.98 |
| O | NM_007218.4 | RNF139 | chr8 | chr8:124474880–124488618 | 3UTR | chr8:124488010–124488312 | 100 | 72.92 |
| P | NM_024598.4 | USB1 | chr16 | chr16:58001407–58021618 | 3UTR | chr16:58020388–58020867 | 159 | 71.21 |
| Q | NM_032812.9 | PLXDC2 | chr10 | chr10:19816432–20289856 | 3UTR | chr10:20284259–20284585 | 108 | 70.8 |
| R | NM_002726.5 | PREP | chr6 | chr6:105273218–105403082 | 3UTR | chr6:105275584–105275934 | 116 | 70.06 |

**Table 2.**
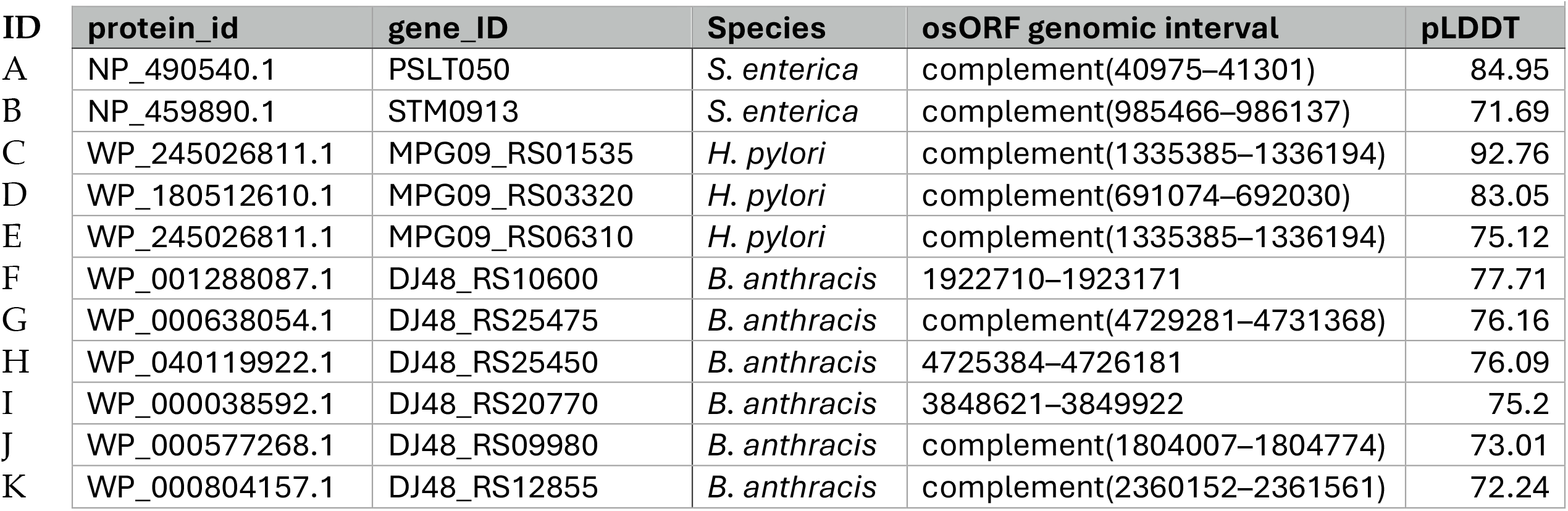
Opposite-strand ORFs within bacterial genes. This table summarizes bacterial osORFs from three different species that yielded an AlphaFold2 / ColabFold pLDDT score of 70 or greater. For each bacterial gene, the protein accession number, gene ID, species, genomic coordinates, and average pLDDT score are reported.

| ID | protein_id | gene_ID | Species | osORF genomic interval | pLDDT |
| --- | --- | --- | --- | --- | --- |
| A | NP_490540.1 | PSLT050 | *S. enterica* | complement(40975–41301) | 84.95 |
| B | NP_459890.1 | STM0913 | *S. enterica* | complement(985466–986137) | 71.69 |
| C | WP_245026811.1 | MPG09_RS01535 | *H. pylori* | complement(1335385–1336194) | 92.76 |
| D | WP_180512610.1 | MPG09_RS03320 | *H. pylori* | complement(691074–692030) | 83.05 |
| E | WP_245026811.1 | MPG09_RS06310 | *H. pylori* | complement(1335385–1336194) | 75.12 |
| F | WP_001288087.1 | DJ48_RS10600 | *B. anthracis* | 1922710–1923171 | 77.71 |
| G | WP_000638054.1 | DJ48_RS25475 | *B. anthracis* | complement(4729281–4731368) | 76.16 |
| H | WP_040119922.1 | DJ48_RS25450 | *B. anthracis* | 4725384–4726181 | 76.09 |
| I | WP_000038592.1 | DJ48_RS20770 | *B. anthracis* | 3848621–3849922 | 75.2 |
| J | WP_000577268.1 | DJ48_RS09980 | *B. anthracis* | complement(1804007–1804774) | 73.01 |
| K | WP_000804157.1 | DJ48_RS12855 | *B. anthracis* | complement(2360152–2361561) | 72.24 |

**Table 3.**
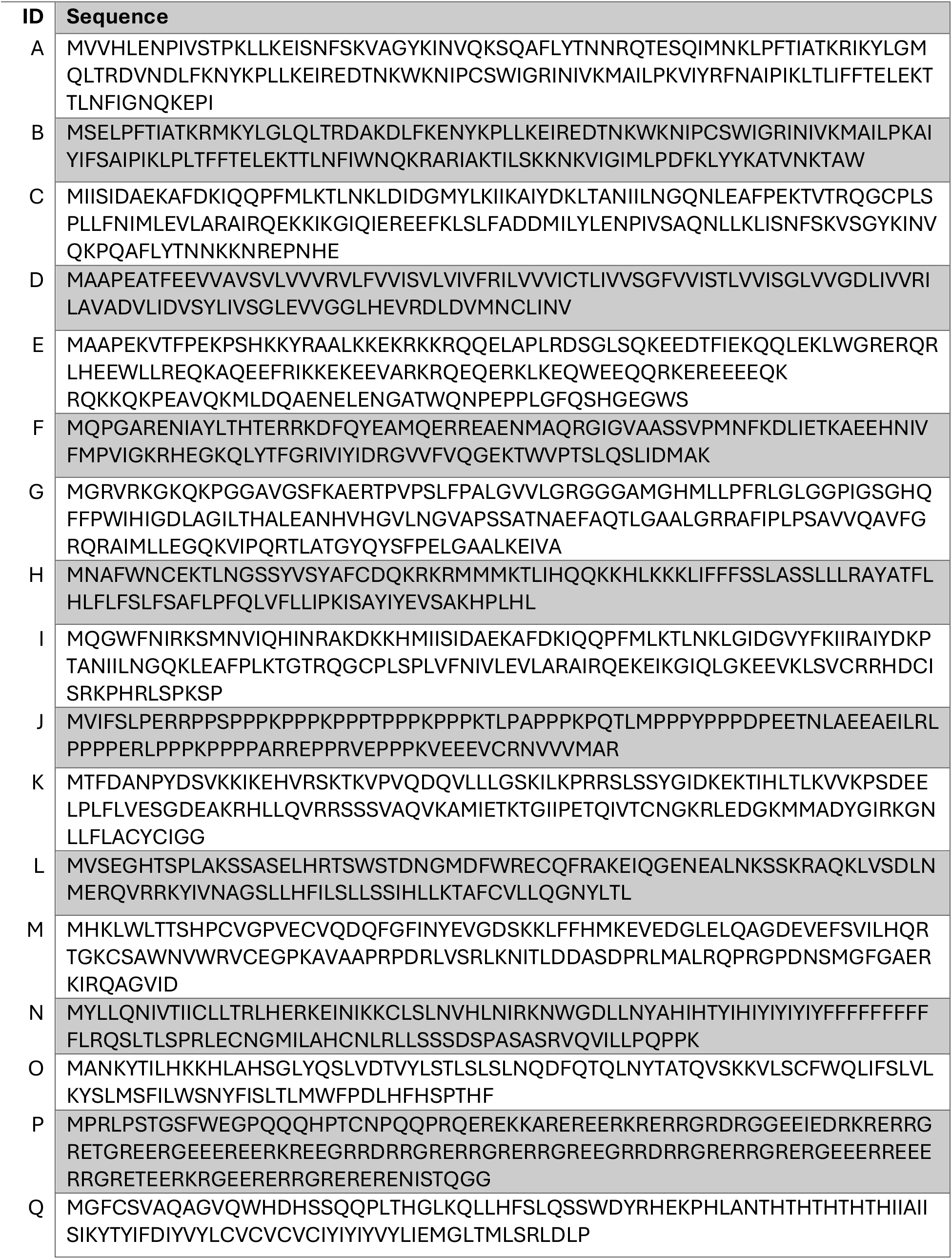
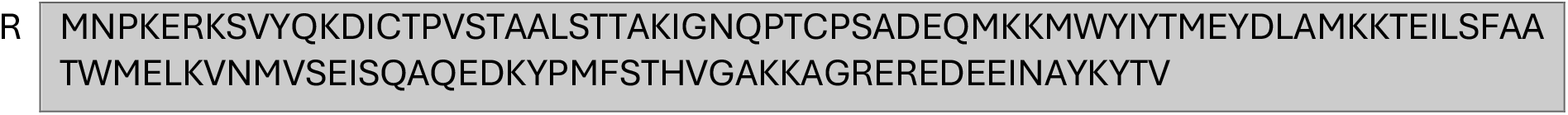
Protein sequences translated from osORFs in Table 1.

**Figure 2.**
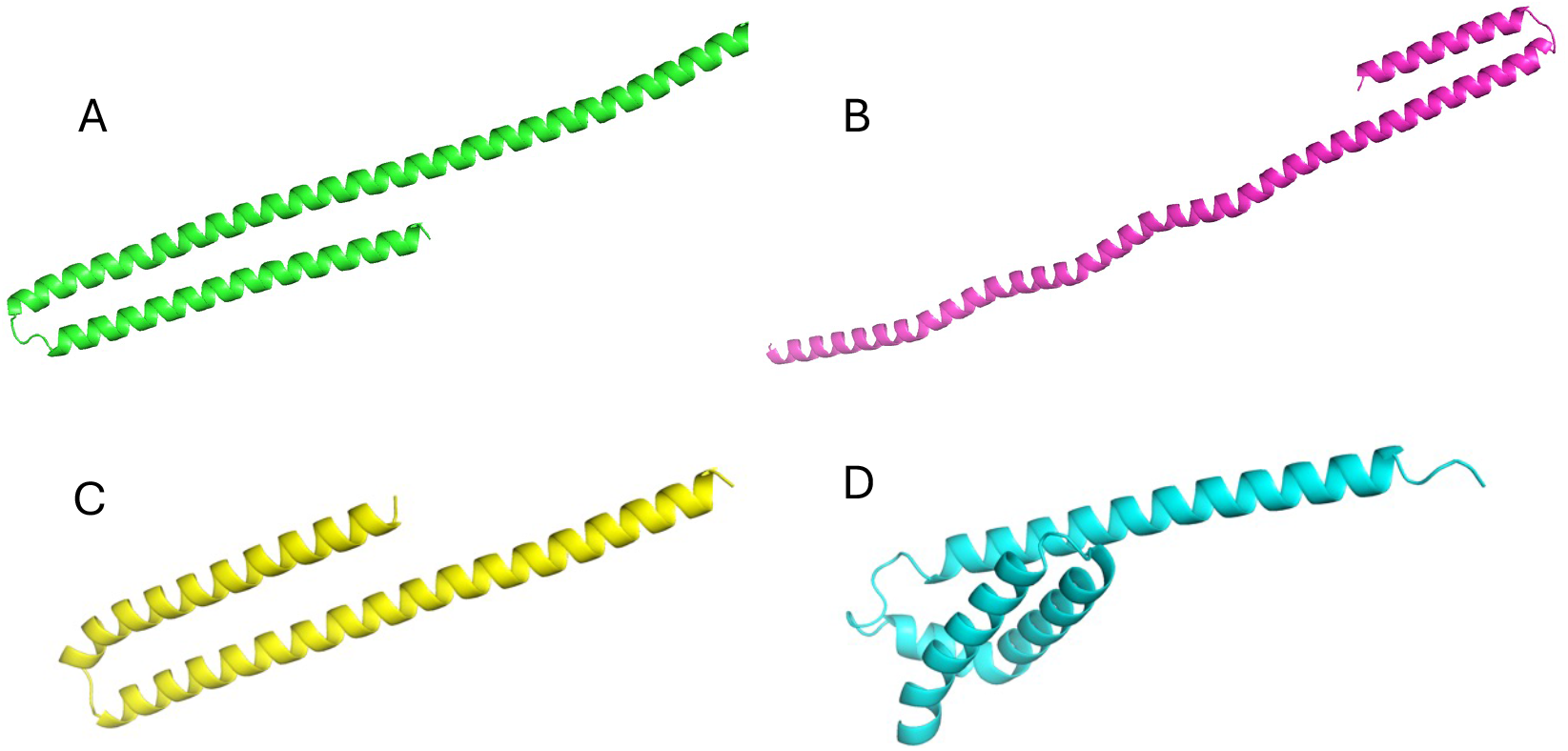
Predicted structures for amino acids sequences translated from opposite-strand ORFs in known bacterial protein-coding genes. (A) 149aa sequence from *B. anthracis* protein WP_040119922.1; (B) 151aa sequence from *B. anthracis* protein WP_000638054.1; (C) 102aa sequence from *B. anthracis* protein WP_000038592.1; (D) 103aa sequence from *H. pylori* protein WP_245026631.1.

This result is consistent with the results reported previously for AntiFam structure predictions [6], where the authors reported AlphaFold2 scores above 80 for six AntiFam sequences that were less than 100aa long. In all six of those cases, the predicted structures were single alpha helices, even simpler than the structures shown in Figure 2 in that the helices did not contain any kinks or turns.

Interestingly, the only bacterial osORF structure scoring above 70 that appeared as something other than a simple helix was a 132aa sequence from an osORF in protein NP_459890 from *S. enterica*, shown in **Figure 3**. In this case, we reviewed the annotation and discovered that NP_459890 was flanked on either side by proteins on the opposite strand, all annotated (including NP_459890) as belonging to a prophage. Because a series of prophage genes should all have the same orientation, NP_459890 appears to be an annotation error, and it should be replaced by the opposite-strand protein whose structure is shown in Figure 3.

**Figure 3.**
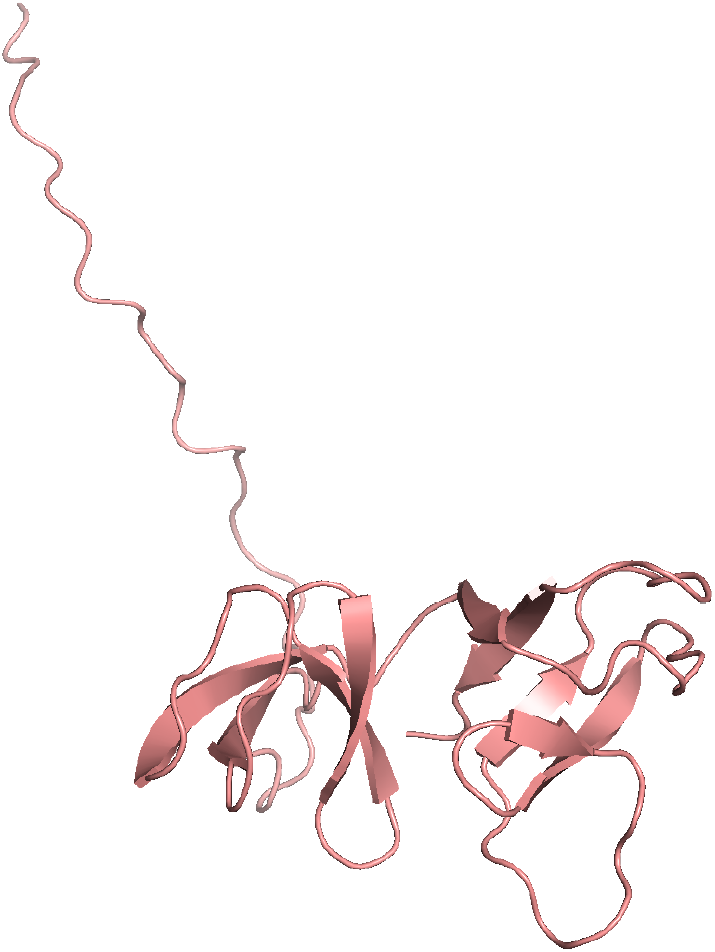
Structure of a protein sequence translated from the reverse strand of *S. enterica* gene NP_459890.1, which received a pLDDT score of 71.7.

Thus of the 1,363 bacterial osORFs >100aa in length, only 11 translated protein sequences scored above 70, and 10 of those are false positives, for an estimated FP rate of 0.73%.

### Human sequences that fold unexpectedly well

One of the more surprising findings was that several of the human osORF translations folded exceptionally well, with six of them scoring above 80 and one scoring 90.47 (**Table 1**). The highest-scoring osORF is a 151aa sequence that appears within NKAIN3 (accession NM_001304533.3), and its predicted structure includes multiple helices and beta sheet sections (**Figure 4**).

**Figure 4.**
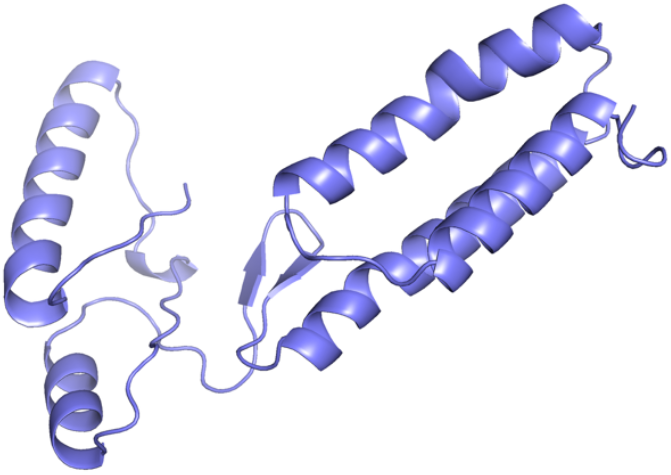
Structure of osORF-NKAIN3, a 151aa protein sequence translated from the opposite strand of NKAIN3, which received a score of 90.47 from ColabFold.

Upon further investigation, we found that NKAIN3 is annotated (in MANE) with an unusually long 3’ UTR, spanning 19,497 bp, which is ten times longer than average for human protein-coding transcripts (the average length for all 3’ UTRs in MANE is 1932 bp).

We then used BLAST [14] to align osORF-NKAIN3 to the NCBI nr database and found multiple significant matches to human and other mammalian proteins. Further analysis revealed that osORF-NKAIN3 is a truncated fragment of the human LINE-1 retrotransposable element ORF2p, a multifunctional enzyme involved in integrating new copies of L1 elements into the host genome [15]. The full ORF2p protein is 1275aa in length; thus the 151aa fragment contained in osORF-NKAIN3 is clearly a nonfunctional copy. We also note that within the NKAIN3 UTR region, the pseudogene spans a larger region than the 151aa initially identified: when we extended the ORF upstream of the start codon, we found homology to ORF2p over a span of 225aa (675bp) ending at position 62,974,148 on chromosome 8.

This result suggests that the annotated 3’ UTR of the MANE isoform of NKAIN3 should be truncated so that it does not include osORF-NKAIN3. Transcriptional evidence from the CHESS annotation [11] places an alternative transcription termination site at position 62,966,300 (**Suppl. Figure S1a**). This proposed site is 18,604bp upstream of the termination site (62,984,904) annotated in MANE and ∼8Kb upstream of the ORF2p pseudogene. The CHESS transcript (CHS.53558.4) has a much shorter 3’ UTR, less than 1 Kb, and osORF-NKAIN3 falls entirely outside the CHESS transcript. Also worth noting here is that although the CHESS isoform encodes a slightly shorter protein sequence, 181aa vs 218aa, its predicted protein structure has a substantially higher pLDDT score than the MANE protein, 78.3 versus 70.0. Thus even though MANE isoforms are generally very high-quality [9], the MANE annotation might be improved by using CHS.53558.4 as the reference transcript for NKAIN3.

Structures for all 18 of the human osORFs with scores >70 are shown in **Suppl. Figure S2**, and their genomic coordinates are in **Table 1**. As the figure shows, 10 of the 18 structures (panels D, E, H, J, L, N, O, P, Q, and R in **Fig. S2**) are simple helices similar to the false structures observed for the bacterial osORFs; these appear to be false positives. However, the other eight human osORFs, including osORF-NKAIN3, show more-complex structures that likely represent pseudogene fragments. We also note that all eight are found within annotated 3’ UTR regions (**Table 1**), and thus do not overlap the protein sequence of the MANE transcript containing them. **Suppl. Figure S1** shows the relative genomic locations of these eight osORFs within the MANE transcripts that contain them.

For three of the eight high-scoring osORFs, in the genes SRRD, KHNYN, and OR2I1P, we observed that the osORF is actually contained within a protein-coding exon of another gene adjacent to and overlapping those genes. In other words, the osORF is actually part of a known protein-coding gene, thus confirming that the ColabFold prediction is a true positive. For these three genes, the genomic position of their 3’ UTR could be adjusted to eliminate the overlaps with adjacent genes. We note that the CHESS database already contains isoforms for SRRD and KHNYN with a shorter 3’ UTR that avoids the overlap.

For the remaining four genes, FRK, ORC4, OR14J1, and ZNF354C, we ran BLAST and confirmed that, as with NKAIN3, their osORFs are pseudogene fragments with very clear homology to other human proteins. In these cases, given that UTRs rarely contain pseudogenes and given that the precise locations of transcription termination sites (TTSs) are difficult to determine [16], the true TTS might well occur before the osORF.

Taken together, the evidence from ColabFold along with sequence homology results for the amino acid translations of these osORFs suggests that the 3’ UTR annotation for these eight human genes should be updated, and that five of the contained osORFs should be annotated as pseudogenes.

## Discussion

To estimate the false positive rate of AlphaFold2 and ColabFold, we generated a set of protein sequences that derived from translations of actual DNA sequences, as opposed to using random or simulated sequences. By choosing open reading frames from the opposite strand of known protein-coding genes, we ensured that the osORFs almost certainly did not represent actual proteins. Because they occur in real genomic sequence, these ORFs might be considered as possible protein-coding genes by automated genome annotation systems.

In total, the structures for 10/1363 bacterial osORFs and 10/7320 human osORFs, all of them at least 300bp in length, yielded pLDDT scores >70. If one used these parameters with AlphaFold2 as a means of identifying likely proteins, then we estimate that 0.23% of predictions (approximately 1/435) based on those structures would be false positives. Note that all of the predictions designated here as false positives created simple structures that consister of a single helix or a helix with one or two kinks in it; thus it requires human intervention to identify these structures. On the other hand, eight out of 18 high-scoring osORFs in the human annotation generated more complex structures that appear to represent true positives, in most cases because they were unannotated pseudogenes. This suggests that in a relatively high proportion of cases (44%), high-scoring structures computed by AlphaFold2 or ColabFold from the translation of genomic regions might identify pseudogenes or other gene-like regions. The identification of these high-scoring amino acid sequences also suggested areas where existing genome annotation could be improved.

## Methods

Genome sequences and annotation were collected for four bacterial genomes: *B. anthracis* strain 2000031021 (GenBank accession GCA_000742655.1), *H. pylori* Hpfe022 (GenBank accession GCA_022925175.1), *S. enterica* subsp. Enterica serovar Typhimurium str. LT2 (GenBank accession GCA_000006945.2), and *S. aureus* strain 199 (GenBank accession GCA_022494545.1). For human annotation, we used release 1.4 of the MANE database [9] as our primary source, and release 3.1.3 of the CHESS database

[11] to search for alternative isoforms as discussed in the main text. We identified all open reading frames on opposite strands of human mature mRNA transcripts; i.e., transcripts with introns spliced out. We similarly found all open reading frames on opposite strands of the bacterial genes, using the span from the start codon to the stop codon to define the search range. For each gene we retained the longest opposite-strand ORF (osORF) if it was at least 300bp long. We required osORFs to start with the canonical start codon ATG and end with any of the three standard stop codons TAA, TGA, or TAG. When multiple ORFs were detected on the reverse strand of a given gene, the single longest ORF was used, ensuring at most one candidate “false protein” per annotated gene. These osORFs were computationally translated into amino acids for input to ColabFold.

### Protein structure prediction and analysis

All sequences were folded with ColabFold, which incorporates the AlphaFold2 pipeline. Each input sequence was processed with the standard single-sequence AlphaFold2 model, and for computational efficiency only one structural model was generated per ORF. All proteins were folded on a local installation of ColabFold running on a single GPU server to ensure consistent computational conditions across all sequences. For each prediction, we extracted the per-residue pLDDT values, which estimate local structural confidence, and from these we computed the mean pLDDT, averaged over all positions in the protein.

Predictions with mean pLDDT ≥70 were classified as high-confidence structures, following the threshold established in [1], which described those models as ones that AlphaFold2 considers generally reliable. Histograms were generated to visualize the distribution of predicted confidence scores for all ORFs. Structures with pLDDT ≥70 were visualized and figures were generated using PyMOL [17].

## Acknowledgements

Thanks to Martin Steinegger for helpful technical discussions on AlphaFold2 and ColabFold. This work was supported in part by NIH grant R01-HG006677.

## Supplementary figure captions

**Supplementary Figure S1.**
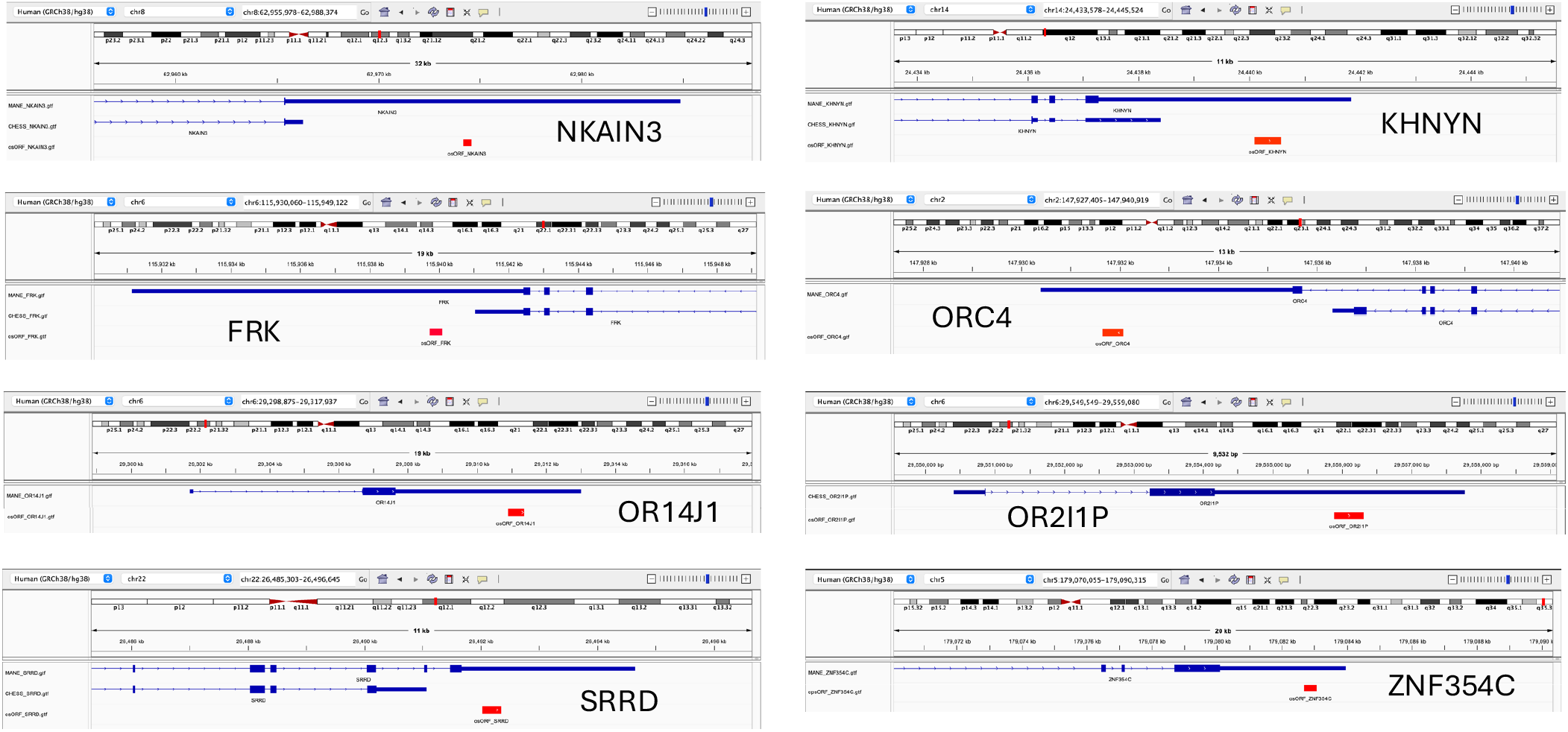
Genomic locations of the opposite-strand open reading frames (osORFs) for the eight osORFs with folding scores of 70 or above as reported by ColabFold. The genes containing the osORFs are, top to bottom and left to right: NKAIN3, FRK, OR14J1, SRRD, KHNYN, ORC4, OR2I1P, and ZNF354C. The exon-intron structure of the gene annotated in MANE is shown in blue across the top, with an alternative isoform with a shorter UTR, taken from the CHESS database, just below it. For OR14J1, OR2I1P, and ZNF354C, the CHESS annotation has the same UTR and is not shown. The position of the osORF is shown as a red rectangle for each gene.

**Supplementary Figure S2.**
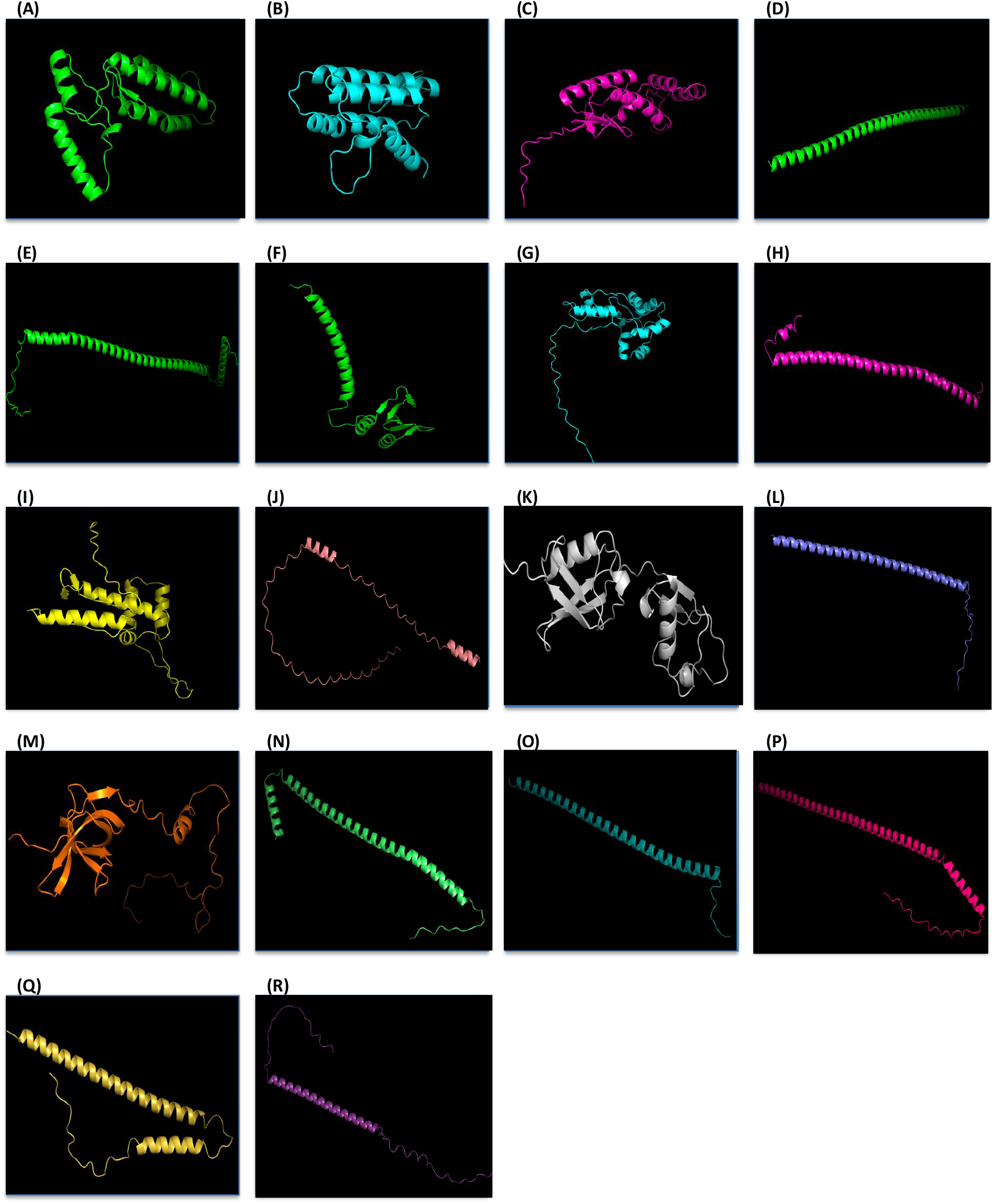
Protein structures predicted by ColabFold for the 18 osORFs in human protein-coding genes with pLDDT scores > 70, as visualized using PyMOL. Letters next to each structure correspond to the osORFs listed in Table 1.

